# Inhibitory CD200-receptor signaling is rewired by type I interferon

**DOI:** 10.1101/2020.02.06.933739

**Authors:** Michiel van der Vlist, M. Inês Pascoal Ramos, Lucas L. van den Hoogen, Sanne Hiddingh, Laura Timmerman, Titus A.P. de Hond, Ellen D. Kaan, Maarten van der Kroef, Robert Jan Lebbink, Florence M.A. Peters, William Khoury-Hanold, Ruth Fritsch-Stork, Timothy Radstake, Linde Meyaard

**Affiliations:** Center for Translational Immunology, Dept. Immunology, University Medical Center Utrecht, Utrecht University, Utrecht, The Netherlands; Oncode Institute, Utrecht, The Netherlands; Center for Translational Immunology, Dept. Rheumatology & Clinical Immunology, University Medical Center Utrecht, Utrecht University, Utrecht, The Netherlands; Sigmund Freud University, Vienna, Austria; Dept of Medical Microbiology, University Medical Center Utrecht, Utrecht University, Utrecht, The Netherlands; Dept of Immunobiology, Yale University School of Medicine, New Haven, CT, The United States of America

**Keywords:** CD200-Receptor, IFNG, Type I Interferon, RasGAP, RASA1, Dok2, SLE, Lupus, Inflammation

## Abstract

CD200 Receptor 1 (CD200R) is an established inhibitory immune receptor that inhibits TLR-induced cytokine production through Dok2 and RasGAP. RasGAP can be cleaved under certain conditions of mild cellular stress. We found that in the presence of cleaved RasGAP, CD200R loses its capacity to inhibit rpS6 phosphorylation. Furthermore, IFNα pre-stimulation of human mononuclear cells results in increased amounts of cleaved RasGAP. Coherently, upon pretreatment with increasing concentrations of IFNα, CD200R gradually shifts from an inhibitor to a potentiator of TLR7/8-induced *IFNG* mRNA production. In peripheral blood mononuclear cells from Systemic Lupus Erythematosus (SLE) patients, a prototypic type I IFN disease, we found an increased proportion of cleaved RasGAP compared to healthy controls. In line with this, in subsets of SLE patients the inhibitory function of CD200R is lost or converted to a potentiating signal for *IFNG* mRNA production. Thus, our data show that type I IFN rewires CD200R signaling and suggest that this cell-extrinsic regulation of signaling could contribute to perpetuation of inflammation in SLE.

## Introduction

Immune inhibitory receptors function to prevent collateral damage by dampening excessive immune cell activation (Ravetch and Lanier, 2000). CD200 Receptor 1 (CD200R) is known as an inhibitory immune receptor that limits pro-inflammatory responses in myeloid cells (Wright et al., 2000). Indeed, CD200fc, a Fc-fusion protein of the best-known ligand for CD200R, has been shown to have immune-suppressive capabilities (Cox et al., 2012; Huang et al., 2018; Simelyte et al., 2008; Yin et al., 2016). Mice deficient in CD200R or in CD200 are healthy under homeostatic conditions. However, upon challenge, these mice generally show increased susceptibility to autoimmune diseases and immune pathology (Hoek et al., 2000; Rygiel et al., 2008; Simelyte et al., 2008; Snelgrove et al., 2008). We previously reported that CD200R inhibits TLR7 signaling, resulting in enhanced anti-viral responses but also enhanced immunopathology upon viral infection in mice (Karnam et al., 2012). All these observations are consistent with an immune-suppressive role for the CD200R-CD200 axis. In line with this, absence of the CD200/CD200R inhibitory axis results in enhanced anti-tumor responses (Kretz-Rommel et al., 2007; McWhirter et al., 2006; Petermann et al., 2007; Rygiel et al., 2012). Coherently, tumor CD200 expression in multiple human cancers is associated with poor prognosis (Aref et al., 2018; Chen et al., 2017; El Din Fouad et al., 2018; Li et al., 2016; Moreaux et al., 2006) and blocking the CD200/CD200R inhibitory axis has been proposed as checkpoint blockade therapy in human cancer (Mahadevan et al., 2019). However, the CD200R-CD200 axis has also been reported as a co-stimulatory axis (Borriello et al., 1997; Wang et al., 2019). Furthermore, CD200R is required to ‘license’ TLR2-mediated immune activation by Herpes Simplex Virus 1 in macrophages (Soberman et al., 2012), and the CD200R/CD200 axis in some cases was suggested to regulate protective anti-tumor responses (Erin et al., 2015; Erin et al., 2018; Liu et al., 2016). Taken together, although CD200R is generally considered to be an inhibitory receptor, there are data that suggest a more versatile and/or complex functionality of CD200R.

CD200R signaling is different from classical inhibitory receptors: CD200R does not signal through an immune tyrosine inhibitory motif (ITIM), but through an NPxY motif (Mihrshahi et al., 2009). CD200R inhibits mitogen activated protein kinases (MAPK) by activating Dok2 and RasGAP/p120 (Mihrshahi et al., 2009; Zhang et al., 2004). The N-terminus of RasGAP/p120 contains the SH2 and SH3 domains that are required to interact with Dok2, the C-terminus contains the GTPase activating protein (GAP) domain to suppress Ras-signaling (Yang et al., 2004). RasGAP/p120 is a target of sub-lethal caspase 3 activity and can be cleaved after amino acid position 455 (Yang et al., 2004). The resulting N-terminal fragment, or ‘fragment N’, has different signaling properties from the full length RasGAP and activates Akt signaling to promote cell survival (Yang et al., 2004). Currently it is unknown what the individual contributions of Dok2 and RasGAP are to CD200R signaling and if cleavage of RasGAP affects the outcome of CD200R ligation.

## Results

### CD200R signaling bifurcates and uses separate pathways to inhibit MAPK and PI3K/Akt

In order to study which signaling pathways are regulated by CD200R we generated U937 cells that ectopically express CD200R-GFP, or a truncated signaling dead control CD200RΔ-GFP (**Fig. S1A**). U937 cells have active Akt, Erk and rpS6 signaling pathways in culture. We assessed the outcome of CD200R ligation on these signaling pathways. Ligation of CD200R-GFP, but not CD200RΔ-GFP with an agonistic antibody (clone OX108) inhibited phosphorylation of mitogen-activated protein kinase (MAPK) Erk at Thr402/Tyr204 (p-Erk) and IL-8 secretion, similar to what was previously reported (**Fig. 1A and S1B**) (Mihrshahi et al., 2009). Additionally, CD200R ligation inhibited phosphorylation of ribosomal protein S6 at Ser234/235 (p-rpS6) (**Fig. 1B**), which is a target of the Erk/MAPK pathway and of the Akt-mTORC1 pathways. Consistently, the inhibitory function of CD200R was not limited to MAPK. We found that CD200R also inhibited Akt at residues phosphorylated by mTORC2 (p-Akt473) and PI3K/PDK (p-Akt308) (**Figs. 1C and 1D**). Similar to antibody-mediated ligation, ligation of CD200R with recombinant human CD200 (rhCD200) inhibited p-Erk in U937 cells (**Fig. S1C**). We conclude that CD200R inhibits both the MAPK/Erk pathway and the Akt pathway and their common downstream target rpS6.

**Figure 1.**
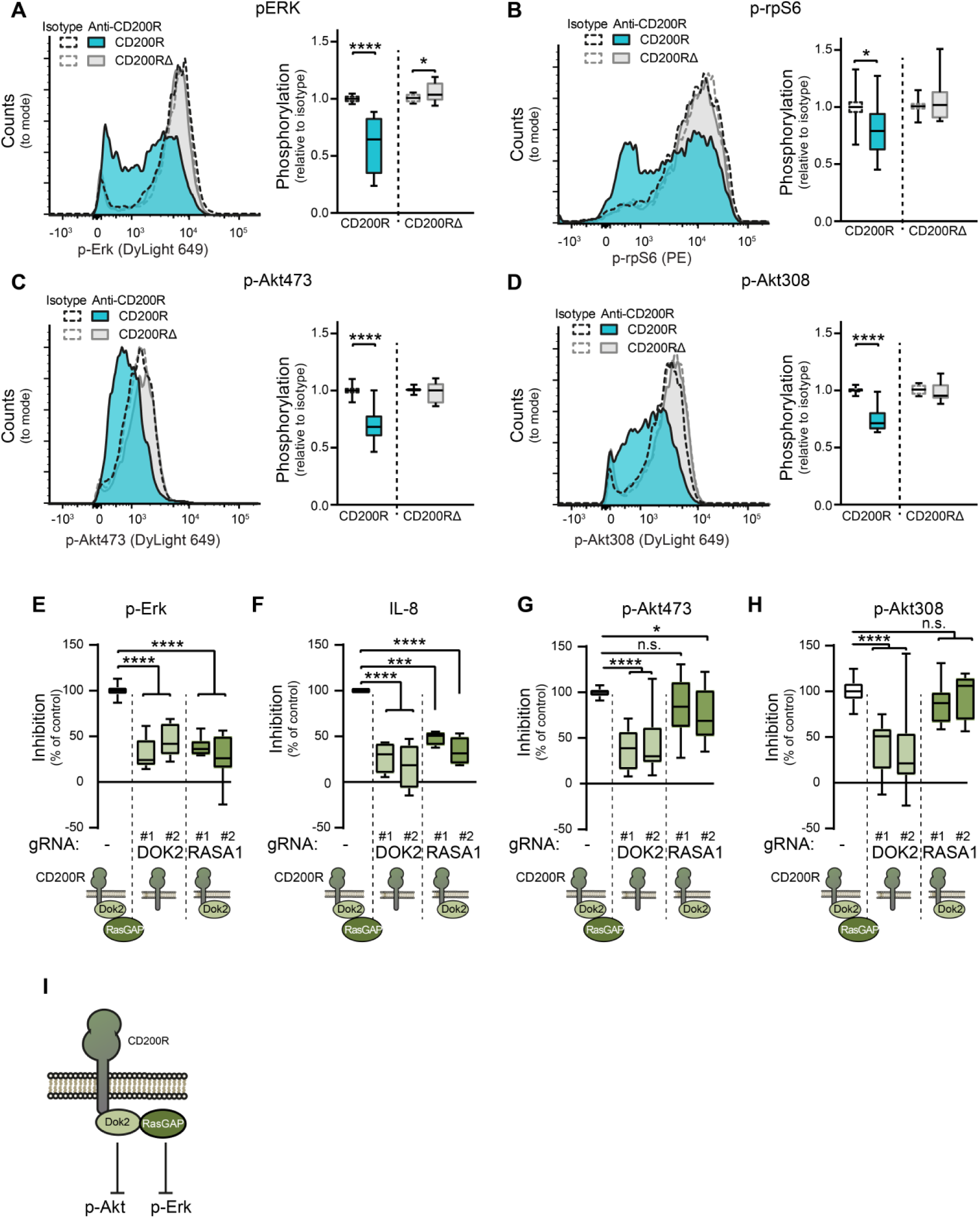
CD200R signaling bifurcates at the level of Dok2. **(A-D)** Phosphorylation of pErk (n=16, from 8 experiments), p-Akt473 (n=14, from 7 experiments), p-Akt308 (n=14, from 7 experiments) and p-rpS6 (n=7, from 7 experiments) in U937-CD200R-GFP (Blue) or CD200RΔ-GFP (Gray) cells after 1h stimulation with anti-CD200R (filled histogram) or isotype control (open histogram) as determined by intracellular flow cytometry. Significance was determined by a t test with Welch correction. The data are normalized to isotype stimulated cells within experiments. See supplemental figure 1A for expression of CD200R and CD200RΔ and gating strategy. (**E-H**) The effect of removal of Dok2 or RasGAP, using two different guide RNA per target, on the inhibitory capacity of CD200R-GFP on (**E**) p-Erk (n=8, from 4 experiments), (**F**) IL-8 secretion (n=4, from 4 experiments), (**G**) p-Akt473 (n=13, from 7 experiments) and (**H**) p-Akt308 (n=13, from 7 experiments). CD200R was ligated with an agonistic antibody or isotype control on U937-CD200R-GFP polyclonal lines deficient for Dok2 or RasGAP. Within experiments, data were normalized to the parental cell line and tested for significance by one-way ANOVA with Holm-Sidak’s correction. Data are obtained in at least 4 independent experiments. Boxplots: median with 25 and 75 percentiles and bars represent minimum and maximum values. (**I**) A model of CD200R signaling. CD200R-signaling bifurcates at the level of Dok2. Dok2 is required for inhibition of Akt and recruitment of RasGAP. Subsequently, RasGAP inhibits Ras/Erk signaling.

CD200R recruits and activates RasGAP through Dok2 (Mihrshahi et al., 2009). To determine the relative contribution of RasGAP and Dok2 in CD200R signaling, we generated U937-CD200R-GFP lines deficient for RasGAP or Dok2 using CRISPR/Cas9 (**Fig. S1D**). We used two different guides that yielded two different deficient U937 cell lines per target, and used these cells as separate polyclonal lines for experiments. In line with previous reports showing that Dok2 is essential for CD200R-mediated inhibition in U937 cells (Mihrshahi et al., 2009), Dok2^-/-^ U937-CD200R-GFP cells failed to inhibit both p-Akt and p-Erk, and IL-8 secretion (**Figs. 1E-H**). In contrast, without RasGAP, CD200R remained able to inhibit p-Akt, while CD200R still failed to inhibit IL-8 secretion and p-Erk (**Figs. 1E-H**). These data show that CD200R-signaling bifurcates at the level of Dok2 and RasGAP, and that in the absence of RasGAP, CD200R still inhibits Akt through a pathway that involves Dok2 (**Fig. 1I**).

### CD200R signaling changes in the presence of cleaved RasGAP

RasGAP is synthesized as a protein of approximately 120kDa, but can be cleaved by caspases during cell stress which disassociates the N-terminal Dok2-binding SH2-domain from the functional C-terminal GAP-domain (Yang et al., 2004). We ectopically expressed a N-terminal fragment of RasGAP that lacks the functional GAP-domain (RasGAP_1-455_) in U937-CD200R cells (**Fig. S2A**) to test the functional consequence of cleaved RasGAP on CD200R function. Overexpression of RasGAP_1-455_ did not affect the amount of p-Erk, p-Akt308, p-Akt473 or p-rpS6 without CD200R ligation (**Fig. S2B**). RasGAP_1-455_ overexpression also did not significantly affect the ability of CD200R to inhibit p-Erk or p-Akt (**Fig. 2A and S2C**). However, CD200R was not able to inhibit p-S6 in presence of RasGAP_1-455_ overexpression (**Fig. 2A**), showing that cleaved RasGAP affects the outcome of CD200R signaling.

**Figure 2.**
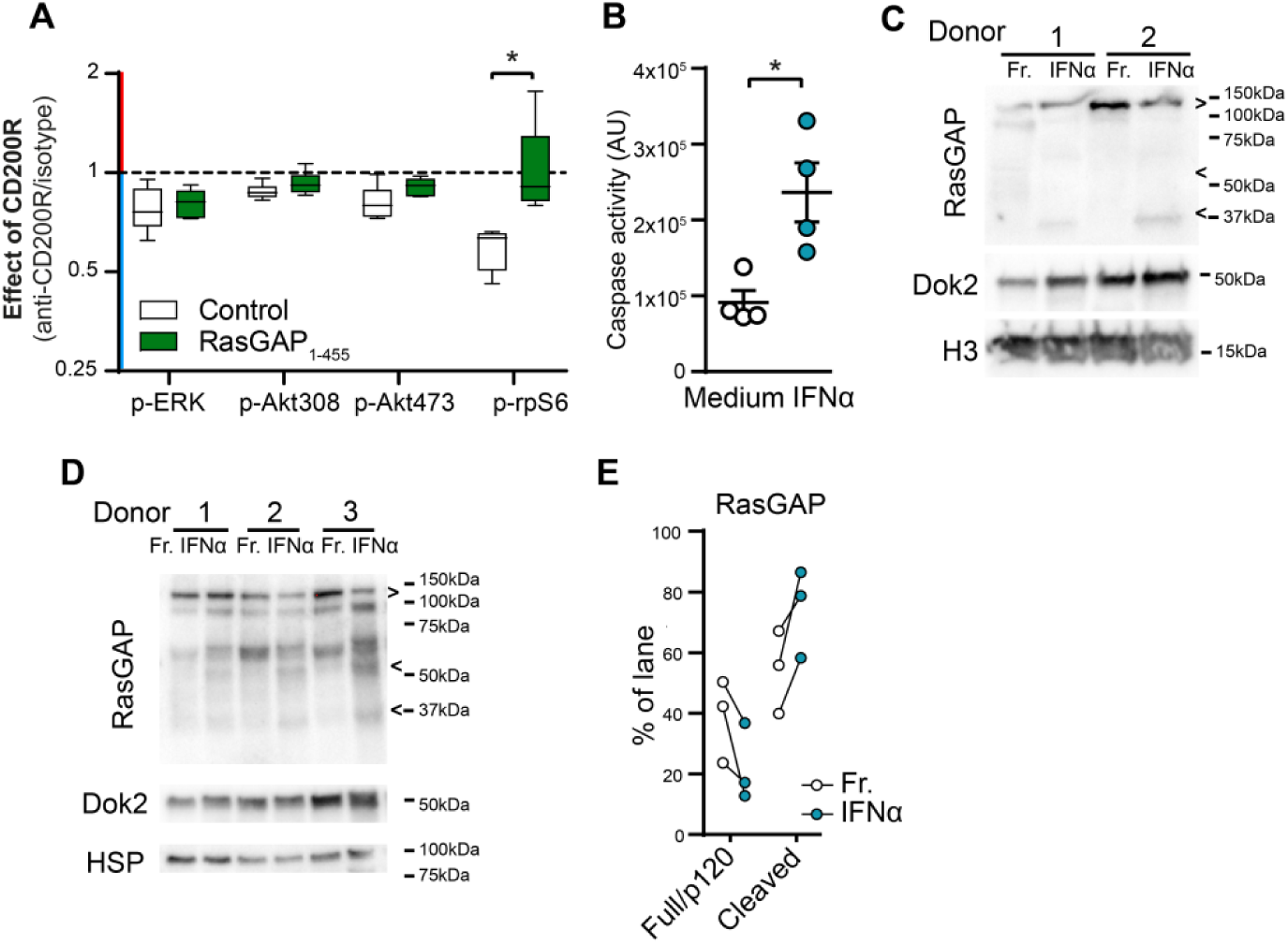
Cleaved RasGAP alters CD200R-signaling. (**A**) Phosphorylation of p-Akt308, p-Akt473, p-Erk, and p-S6 after 1h stimulation of CD200R with agonistic-antibodies or isotype control in CD200R-U937-cells that over-express full-length RasGAP/p120 or RasGAP_1-455_. Data were normalized to isotype-stimulated cells. Significance was determined by a t test with Welch correction (n=6). (**B**) Caspase activity in fresh and IFNα stimulated PBMC as measured by CaspGlo 3/7 assay. Significance was determined by a t test (n=4). (**C, D**) Western blot analysis of RasGAP and Dok-2 protein expression and cleavage in fresh (Fr.) or 16h IFNα stimulated cells. Blots are lysates of (C) CD14^+^ monocytes and (D) PBMC cultures. >: full length RasGAP/p120; <: published RasGAP cleavage fragment (Yang et al., 2004). H3: Histone 3. HSP: heat shock protein 90. (**E**) Quantification of Figure 2D. Data represent the amount full length RasGAP/p120 or the amount of cleaved RasGAP as percentage of signal in the whole lane. Significance was determined by paired-t test.

### Type I IFN exposure induces RasGAP cleavage

RasGAP can be cleaved by caspase 3 (Yang et al., 2004). In human healthy control peripheral blood mononuclear cells (PBMCs) we found that type I IFN can activate caspase 3 activity (**Fig. 2B**), and thus potentially affects RasGAP cleavage. Indeed, 16h exposure to type I IFN (IFNα) increased the amount of cleaved RasGAP fragments in PBMC and isolated monocytes (**Figs. 2C-2E**). We found the previously reported RasGAP cleavage fragments (fragment N of ∼56kDa and fragment N2 of ∼40kDa (Yang et al., 2004)), but also products of other sizes suggesting the involvement of other proteases. As opposed to RasGAP, IFNα pre-stimulation of PBMC and monocytes did not noticeably affect Dok2 protein expression (**Figs. 2C-2E**). Thus, IFNα stimulation induces RasGAP cleavage which may lead to a change in CD200R function.

### Type I IFN exposure converts CD200R from an inhibitory to immunostimulatory receptor

In absence of RasGAP, CD200R selectively inhibited Akt (Fig. 1). PI3K/Akt can provide negative feedback on the production of type I cytokines (Fukao et al., 2002; Ruhland and Kima, 2009). To clarify the role of Akt and Erk in the regulation of TLR7/8 signaling, we used selective inhibitors of Akt (AKTVIII, 2μM) and Erk (U0126, 2.5μM) in primary human monocyte derived dendritic cells (moDCs) that endogenously express functional CD200R (**Figs. S2C and S2D**), stimulated with the TLR7/8 ligand R848. While inhibition of Erk resulted in reduced secretion of IL-8, inhibition of Akt potentiated secretion IL-8 and IL-1β in the majority of the donors (**Fig. 3A**). This demonstrates that selective inhibition of Akt, which occurs if only Dok2 and not RasGAP/p120 is functionally recruited to CD200R, could potentiate TLR7/8 mediated responses. Therefore, we assessed the function of CD200R in freshly replated moDCs versus 16h IFNα pre-exposed moDCs seeded on immobilized recombinant human CD200 (rhCD200). In freshly replated cells, CD200R ligation by CD200 inhibited TLR7/8 ligand R848-induced p-Akt473 and p-rpS6 (**Fig. 3B, 3C and S2E**). IFNα pre-exposure marginally affected the ability of CD200R to inhibit R848-induced p-Akt473 (**Fig. 3B**). However, after pre-exposure to IFNα, CD200R lost inhibitory capacity for p-rpS6 in all donors and even potentiated R848-induced p-rpS6 in some donors (**Fig. 3C**). Overall, we conclude that IFNα driven inflammation favors production of cleaved RasGAP fragments which re-wires the CD200R signaling cascade.

**Figure 3.**
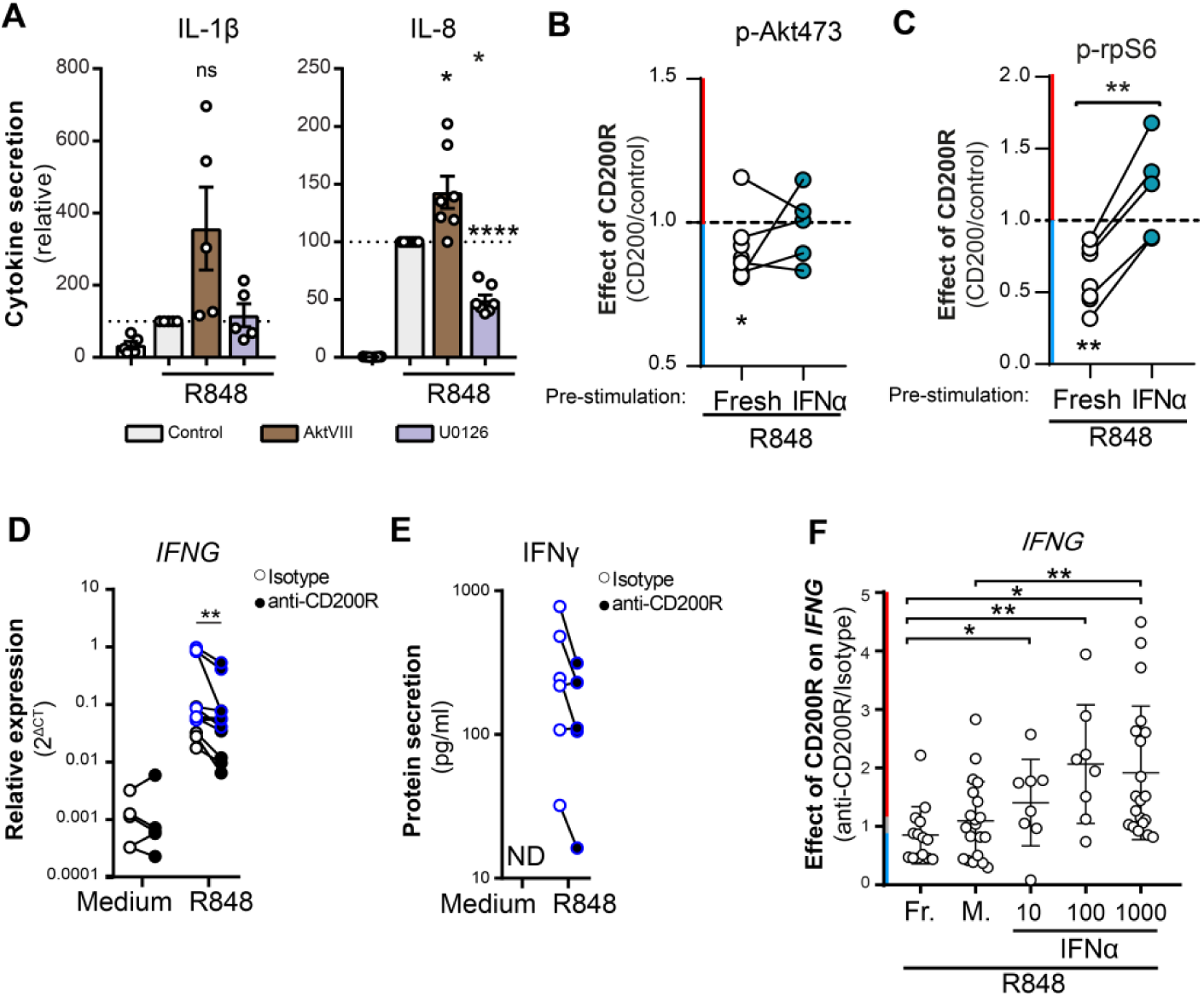
Exposure to type I IFN reverses CD200R function in PBMC. (**A**) Cytokine secretion by moDC stimulated with R848 for 6h in presence of control, Akt-inhibitor AktVIII or Mek-inhibitor U0126. IL-1β: n=5, IL-8: n=8. Significance was determined by one-sample t test. (**B, C**) Phosphorylation of p-Akt473 and p-rpS6 in HC moDC. Fresh replated (n=8, 3 samples without IFN-counterpart) or IFNα-stimulated (16h, n=5, paired with fresh samples) moDCs were seeded on plates coated with control (recombinant human CD200R1, rhCD200R1) or recombinant human CD200 (rhCD200). After 2h pre-incubation, the moDC were stimulated with 1µg/ml R848 for 30 minutes. Phosphorylation was assessed by flow cytometry. Depicted is the ratio of phosphorylation in CD200-stimulated conditions over control stimulated conditions. Significance between the effect of CD200R on fresh and IFNα stimulated cells was determined by t test. Significance of the effect of CD200R in medium or IFNα stimulated cells was determined by one-sample t test. (**D, E**) mRNA (n=11) and protein (n=6) data on R848-induced IFNG or IFNγ production in PBMC. Donors with blue lines are matched between mRNA and protein data. Significance was determined by Wilcoxon test. ND = not detectable. (**F**) The ratio of R848 induced *IFNG* mRNA production of PBMC stimulated with anti-CD200R over PBMC stimulated with isotype. *IFNG* mRNA production was measured in healthy control PBMC that were used fresh after isolation (Fr., n=14), or have been cultured for 16h in medium (Med, n=20), 10 (n=8), 100 (n=8) or 1000 (n=20) U/ml of IFNα prior to ligation of CD200R with an agonistic antibody and stimulation of TLR7/8 with R848 for 4h. Significance was determined by a paired t test with Welch correction.

We next assessed the inhibitory capacity of CD200R in PBMC from healthy controls (HC). In PBMC cultures, we found that *IFNG* mRNA and IFNγ protein production was induced by R848 and inhibited by CD200R ligation with immobilized antibodies (**Figs. 3D and 3E**). *IFNG* mRNA in these cultures was most likely produced by NK cells, but orchestrated by monocytes (**Figs. S3A-S3C**) because depletion of either NK cells or monocytes(Chan et al., 1991; Okamura et al., 1995), but not T cells, prevented production of *IFNG* mRNA. We tested the effect of IFNα on CD200R function in human PBMC by using freshly isolated cells, or cells that were pre-stimulated for 16h culture in medium with increasing doses of IFNα. Freshly isolated or pre-treated HC PBMC were seeded on immobilized agonistic CD200R antibody followed by stimulation with R848. CD200R ligation inhibited R848-induced *IFNG* mRNA expression in freshly isolated PBMC from most healthy controls.

However, pre-treatment of PBMC with IFNα abrogated or reversed the capacity of CD200R to inhibit *IFNG* mRNA in almost all healthy donors (**Figs. 3F and S3D**). With increasing dose of IFNα, the ratio of *IFNG* mRNA production by CD200R stimulated cells over isotype control antibody stimulated cells, changed from <1 (inhibition) to 0 (no effect of CD200R) to >1 (potentiation by CD200R). Of note, after overnight culture of PBMC, the inhibitory function of CD200R was also partially lost compared to freshly isolated cells (**Fig. 3F**). This coincided with increased mRNA expression of type I IFN-stimulated genes *IFITM1* and *MX1*, suggesting a type I IFN-response induced by overnight culture **(Fig. S3E)**.

### Increased RasGAP cleavage in systemic lupus erythematosus PBMC

Type I IFN is a feature of many systemic auto immune diseases and its pathogenic role is well established for systemic lupus erythematosus (SLE) (Crow and Rönnblom, 2018). Blocking the type I interferon receptor subunit 1suppresses disease activity in SLE patients (Morand et al., 2020). SLE predominantly affects women (9:1 ratio) (Bennett et al., 2003) and next to type I IFN, TLR7 is implicated in its pathogenesis (Souyris et al., 2018). Furthermore, caspase activity is upregulated in SLE PBMC (Krishnan et al., 2005). We therefore assessed the availability of Dok2 and RasGAP in PBMC of SLE patients. At the protein level, both Dok2 and RasGAP expression is slightly decreased in PBMC from SLE patients (**Fig. 4A**). However, similar to what we observe after IFNα pre-stimulation in healthy control PBMC, we found an increase in cleaved RasGAP fragments in PBMC from SLE patients (**Fig. 4A**). This results in a lower ratio of RasGAP/p120 over cleaved RasGAP (**Fig. 4B**). Together these data suggest that RasGAP/p120 cleavage is enhanced by IFNα *in vivo*, which is exemplified in SLE PBMC.

**Figure 4.**
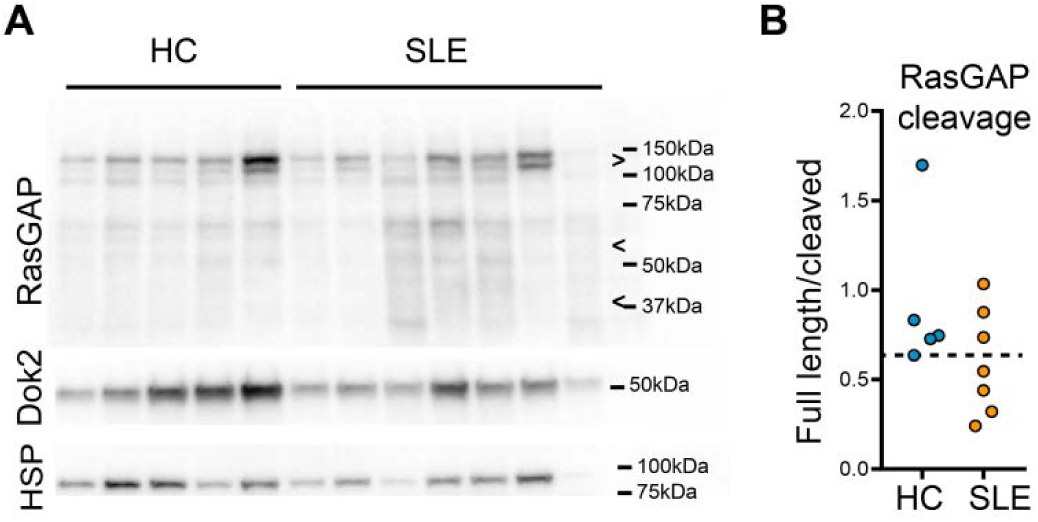
Cleaved RasGAP in systemic lupus erythematosus patients. (**A**) Western blot analysis of RasGAP and Dok-2 protein expression in healthy control (HC) or systemic lupus erythematosus (SLE) patients’ PBMC. >: Full length RasGAP/p120, <: known caspase 3 cleavage fragments (Yang et al., 2004). HSP: heat shock protein 90 loading control. (**B**) Quantification of Figure 4A. Data represent the ratio of full length RasGAP/p120 or the amount of cleaved RasGAP as percentage of signal in the whole lane. Significance was determined by Wilcoxon test, p=.2677.

### CD200R function is abrogated or inverted in PBMC from SLE patients

The observed increased RasGAP cleavage could result in rewiring of the CD200R signaling in (part of) SLE patients. To test this, we used PBMC from clinically well-defined SLE patients and age- and sex-matched healthy controls (HC) in two different cohorts. Antibody-mediated CD200R ligation inhibited TLR7/8-mediated *IFNG* mRNA expression in most HC PBMC in cohort 1 (**Fig. 5A, Table S1**). In contrast, in SLE PBMC with a detectable *IFNG* mRNA response, with the exception of one patient, CD200R was unable to inhibit *IFNG* expression, and even potentiated *IFNG* expression in response to TLR7/8 ligation in 5 out of 7 SLE patients (**Fig. 5A, Table S1**). In a separate replication second cohort, CD200R function was also lost or inverted in 7 out of 11 SLE patients (**Fig. 5B, Table S1**). We excluded that CD200R function was lost because of lower CD200R expression: instead in SLE patients CD200R expression was increased on non-classical inflammatory CD14^-^CD16^+^ monocytes (**Fig. S4A)**, and on CD56^+^ NK cells and CD19^+^ B cells (**Fig. S4B**). Medication used for SLE patients could potentially affect TLR7/8 responses and therefore CD200R function. Indeed, patients treated with hydroxychloroquine (HCQ), prednisone (Pred) or azathioprine (Aza) had decreased *IFNG* mRNA production after R848 stimulation (**Fig. S4C**). However, none of these drugs had an influence of the function of CD200R in these patients (**Fig. S4C**). Coherently, HCQ pre-treatment of HC PBMC did not change the function of CD200R (**Fig. S4D**), confirming that the functional switch we observed for CD200R was not the result of medication. Interestingly, the reversal of CD200R function was most prominent in SLE patients with Lupus Nephritis (LN), a severe manifestation of SLE, suggesting a link between aberrant CD200R inhibition and disease severity (**Fig. 5C**).

**Figure 5.**
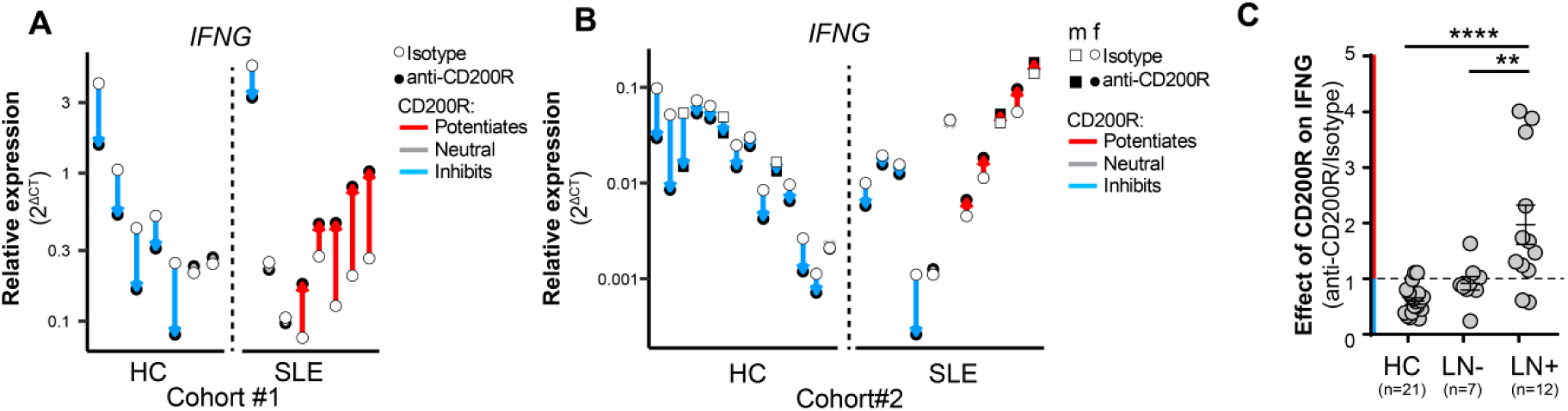
CD200R potentiates IFNG mRNA production in systemic lupus erythematosus patients. (**A, B**) *IFNG* mRNA production relative to *GAPDH* (A) or *B2M* (B) by healthy control and SLE patients’ PBMC in two cohorts. PBMC were seeded on an isotype control (open circles) or an agonistic anti-CD200R antibody (filled circles) prior to stimulation with R848 for 4h. Each connected dot represents one patient/control. The color of the connecting arrow indicates: inhibition by CD200R ligation (blue, >15% inhibition compared to isotype control); potentiation by CD200R ligation (red, >15% potentiation compared to isotype control); or no effect of CD200R ligation (grey, <15 % inhibition or <15% potentiation compared to isotype control). (**C**) Ratio of R848-induced *IFNG* mRNA production from data in fig. 5A and 5B, grouped to HC, SLE patients without lupus nephritis (LN-) and patients with (a history of) lupus nephritis (LN+). Significance was determined with a one-way ANOVA.

## Discussion

Our data show that CD200R signaling bifurcates at the level of Dok2, inhibiting the PI3K/Akt pathway through Dok2, and the Ras/Erk pathway through recruitment of both RasGAP to Dok2. Type I IFN exposure results in RasGAP cleavage which changes the outcome of CD200R signaling. Consequently, CD200R potentiates R848-induced *IFNG* mRNA production in PBMC from HC that are exposed to type I IFN, and in untreated PBMC of part of SLE patients. This functional switch is not binary: the functional outcome of CD200R ligation seems to progress along a sliding scale from inhibition to potentiation. Previously, immune receptors with an immune tyrosine switch motif (ITSM) were shown to relay an activating or inhibitory signal by using different adaptor molecules (Wu and Veillette, 2016). For CD200R, we now propose a different mechanism, where extracellular cues can rewire the intracellular signaling of an immune receptor by inducing cleavage of their adaptor molecules. Hence, CD200R is a versatile receptor that, depending on cell-extrinsic cues, can function anywhere between an inhibitory to potentiating immune receptor.

How would the immune system benefit from this versatility of CD200R? In homeostatic conditions, ligation of CD200R results in suppression of TLR responses in healthy donors (**Fig. S5A**). This is in line with published work in CD200 or CD200R knock-out mice showing stronger inflammatory responses and immunopathology (Hoek et al., 2000; Karnam et al., 2012; Rygiel et al., 2009; Snelgrove et al., 2008), and that treatment with CD200-fc prior to disease onset dampens renal pathology in lupus prone mice (Yin et al., 2016). We reason that this inhibitory function is a useful threshold for TLR-mediated immune activation and keeps immune cells quiescent via CD200 expressed on, for example, endothelial cells, glomeruli of the kidney, and apoptotic cells (Li et al., 2012; Rosenblum et al., 2004; Wright et al., 2001).

However, we hypothesize that in the context of infection, immune cells would need to induce rather than inhibit type I immunity to clear intracellular pathogens (**Fig. S5B**). Our data show that in such an inflammatory milieu, CD200R can switch function to potentiate TLR-induced *IFNG* production, a hallmark type I immunity cytokine. This potentially enhances intra-cellular pathogen clearance, after which inflammation will cease. Thus, we hypothesize that the versatility of CD200R aids the immune system to quickly adapt to changing inflammatory environments. This beneficial feature of CD200R goes awry the moment there is type I IFN production in the absence of an (infectious) source of inflammation that can be cleared, as is the case for SLE (**Fig. S5C**). In a subset of SLE patients, CD200R indeed potentiated *IFNG* mRNA production.

We found that CD200R signaling impinges on Erk, Akt and their common downstream target rpS6. These targets can explain potentiation of R848-induced *IFNG* at several levels. Akt can provide negative feedback on TLR signaling (Fukao et al., 2002; Ruhland and Kima, 2009). Indeed, inhibition of Akt potentiated inflammatory cytokine secretion in human moDC. Our data show that CD200R selectively inhibits Akt in absence of RasGAP, which would inhibit a negative feedback loop which leads to increased *IFNG* production. Additionally, in presence of cleaved RasGAP and after type I IFN treatment, CD200R loses its inhibitory capacity towards rpS6, a target of mTORC1. mTORC1 is involved in inflammasome activation (Moon et al., 2015), which via IL-18 production can drive *IFNG* mRNA production (Chan et al., 1991; Okamura et al., 1995). Rapamycin, an inhibitor of mTORC1, reduces lupus disease activity, measured among others by SLEDAI score (Fernandez et al., 2006; Lai et al., 2018). Besides direct inhibition of *IFNG* production by T cells, rapamycin could overrule the CD200R-effect on rpS6 as an additive layer for the effect of rapamycin treatment of SLE patients. Overall, the type I IFN-mediated rewiring of CD200R signaling to potentiate rpS6 signaling could contribute to the chronicity of SLE.

Interfering with the CD200-CD200R pathway is suggested to be a therapeutic opportunity in cancer and auto-immunity (Kretz-Rommel and Bowdish, 2008; Mahadevan et al., 2019; Simelyte et al., 2008). However, depending on the setting, the CD200-CD200R axis is reported to either have pro- and anti-tumor effects. Our data show that the outcome of CD200R-ligation depends on the inflammatory milieu. Thus, the outcome of therapeutic targeting of the CD200-CD200R axis will likely depend on the inflammatory status of the patient at the moment of treatment.

In conclusion, we demonstrate that the signal transduction machinery of the immune-inhibitory receptor CD200R is responsive to type I IFNs thereby allowing CD200R to progressively switch from inhibitory to stimulatory signaling depending on the environment. As a general concept, the rewiring of inhibitory receptors by inflammatory cues has implications for therapeutic approaches targeting inhibitory receptors in patients.

## Acknowledgments

We thank all members of our lab for the fruitful discussions; Marion H. Brown (University of Oxford) and Christian Widmann (University of Lausanne) for reagents; Ruslan Medzhitov (Yale University) and Hans Clevers (Hubrecht Institute) for the conceptual discussions; and Louis Bont (UMC Utrecht) and Klaas P.J.M. van Gisbergen (Sanquin) for their critical comments on the manuscript.

## Funding

MV and LM are supported by the Netherlands Organization for Scientific Research (NWO), (ALW Grants 863.14.016, 821.02.025 and NWO Vici 918.15.608). FMAP, MIPR, SH and EDK are supported by the Dutch Arthritis Foundation (Grants 06-1-403, 12-2-101, 14-02-201, 18-1-403).

## Author contributions

Conceptualization (MV, MIPR, TR, LM), Methodology (MV, MIPR, TR, LM), Formal Analysis (MV, MIPR, LH, SH), Investigation (MV, MIPR, LH, SH, MK, FMAP, LT, TAPH, EDK), Resources (RJL), Patient inclusion (LH, RF, TR), Writing original draft (MV), Writing reviewing and editing (LM, TR, WKH, MIPR), Visualization (MV, MIPR), Supervision (MV, TR, LM), Project Administration (MV, LM), Funding Acquisition (MV, LM).

## Declaration of Interests

The authors declare no competing interests.

## Materials and Methods

### Antibodies

| Target | Label | Vendor | Clone | Catalog# | usage | RRID |
| --- | --- | --- | --- | --- | --- | --- |
| CD200R1 | Unlabeled | eBioscience | OX108 | 16-9201-85 | 10ug/ml | AB_10804889 |
| Isotype control IgG1 | Unlabeled | eBioscience | P3.6.2.8.1 | 16-4714-85 | 10ug/ml | AB_470162 |
| CD200R1 | Unlabeled | In house | OX108 | - | 3ug/ml | n.a. |
| SIRL1/isotype | Unlabeled | In house | 1A5 | - | 1ug/ml | n.a. |
| Akt p-Thr308 | Unlabeled | CST | D25E6 | 13038 | 3600x | AB_2629447 |
| Akt p-Ser473 | Unlabeled | CST | D9E | 4060 | 200x | AB_2315049 |
| Erk pThr402/Tyr204 | Unlabeled | CST | D13.14.4E | 4370 | 200x | AB_2315112 |
| rpS6 pSer234/235 | PE | CST | D57.2.2E | 5316 | 200x | AB_10694989 |
| CD14 | APC-H7 | BD Biosciences | MφP9 | 560180 | 50x | AB_1645464 |
| CD14 | APC-eF780 | eBioscience | 61D3 | 47-0149-42 | 100x | AB_1834358 |
| CD16 | V500 | BD Biosciences | 3G8 | 561394 | 50x | AB_10611857 |
| CD19 | PerCP | Biolegend | HIB19 | 302228 | 100x | AB_893272 |
| CD19 | AF700 | eBioscience | HIB19 | 56-0199-41 | 50x | AB_2043820 |
| CD200 | APC | eBioscience | OX104 | 17-9200 | 10x | AB_2573297 |
| CD200R | PE | eBioscience | OX108 | 12 9201 | 20x | AB_1210851 |
| CD3 | AF700 | Biolegend | HIT-3a | 300324 | 40x | AB_493739 |
| CD3 | PerCP | Biolegend | UCHT1 | 300428 | 20x | AB_893298 |
| CD56 | FITC | Biolegend | HCD56 | 318304 | 20x | AB_604100 |
| CD56 | AF700 | BD Biosciences | B159 | 557919 | 50x | AB_396940 |
| CD56 | PE-CF594 | BD Biosciences | B159 | 562289 | 25x | AB_11152080 |
| HLA DR | FITC | BD Biosciences | M80 | 347363 | 25x | AB_400291 |
| HA | APC | Miltenyi | GG8-1F3.3.1 | 130-098-404 | 50x | AB_2751024 |
| RasGAP | Unlabeled | Santa Cruz | B4F8 | Sc-63 | 1000x | AB_628206 |
| Histon 3 | Unlabeled | CST | Poly clonal | 9715 | 1000x | AB_331563 |
| Dok2 | Unlabeled | Santa Cruz | H192 | sc-13952 | 200x | AB_2095091 |
| Donkey-a-Rabbit | AF647 | ThermoFisher | Poly clonal | A31571 | 400x | AB_162542 |
| Donkey -a -Rabbit | AF594 | ThermoFisher | Poly clonal | A21207 | 400x | AB_141637 |
| Goat-a-Rabbit | DL649 | Jackson | Poly clonal | 111-496-003 | 200x | n.a. |
| Donkey-a-Rabbit | AF488 | ThermoFisher | Poly clonal | A11055 | 400x | AB_2534102 |
| Goat-a-Rabbit | HRP | ThermoFisher | Poly clonal | 31460 | 5000x | AB_228341 |
| Goat-a-Mouse | HRP | Jackson | Poly clonal | 11-005-174 | 5000x | n.a. |

### Gene expression quantification

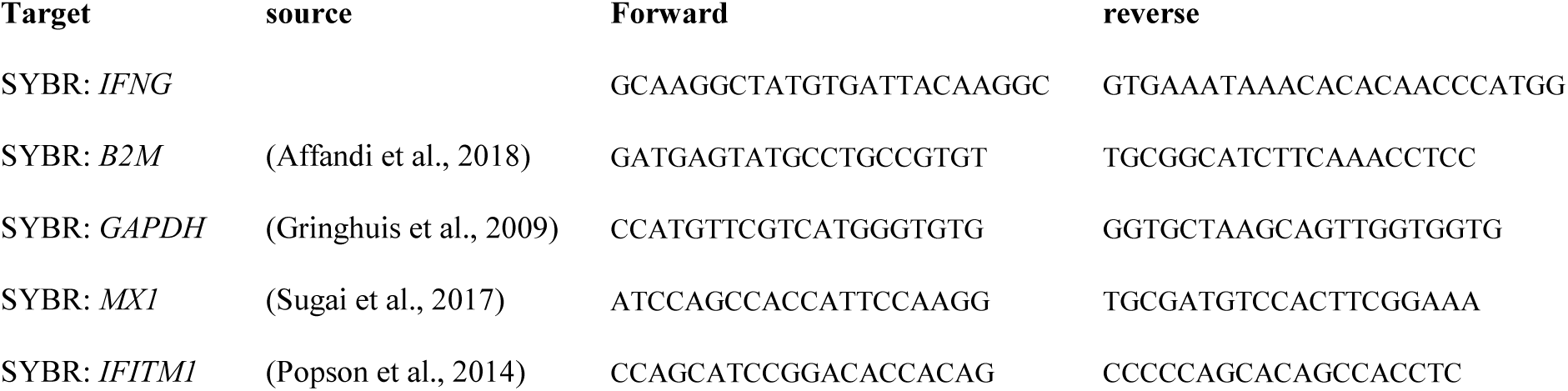

### Stimuli

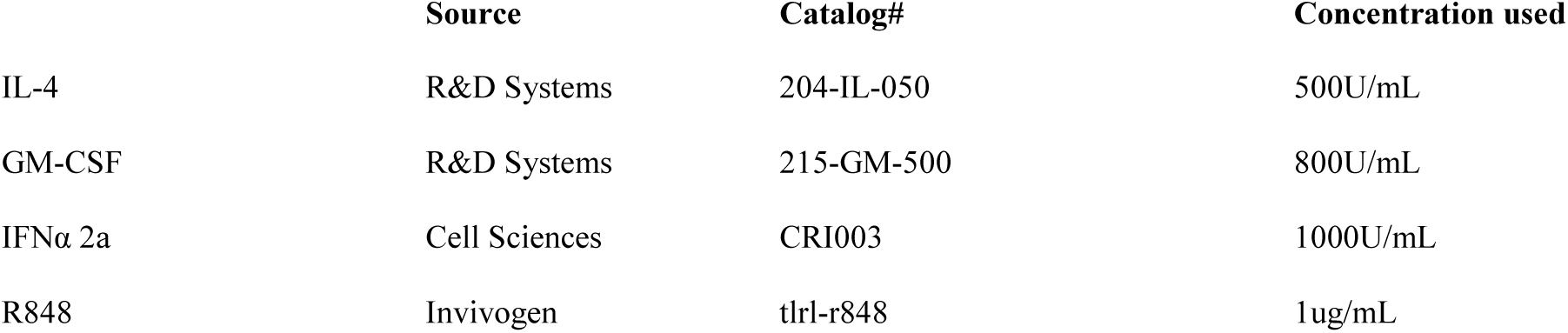

### Recombinant DNA

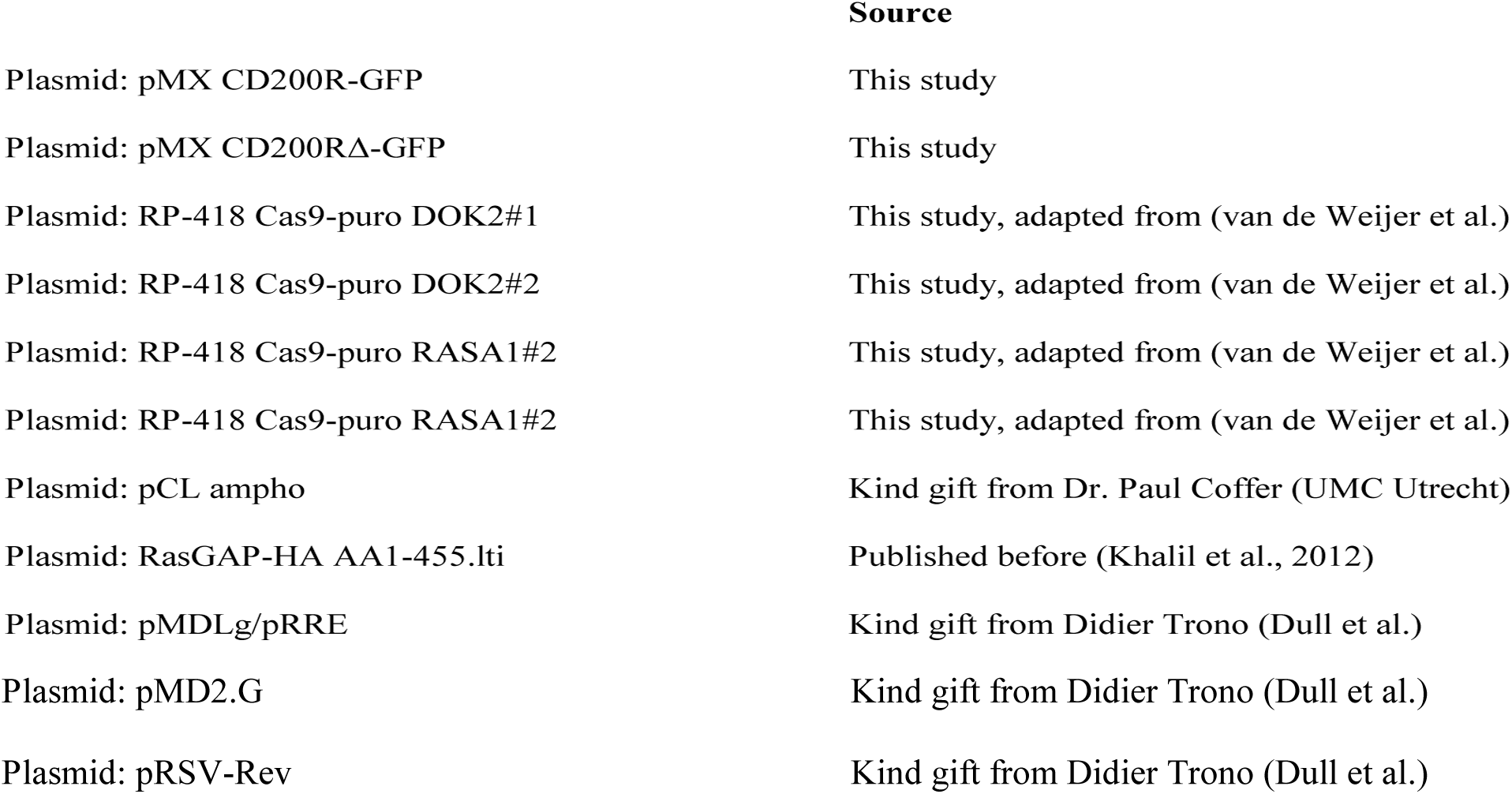

### Experimental Models: Cell Lines

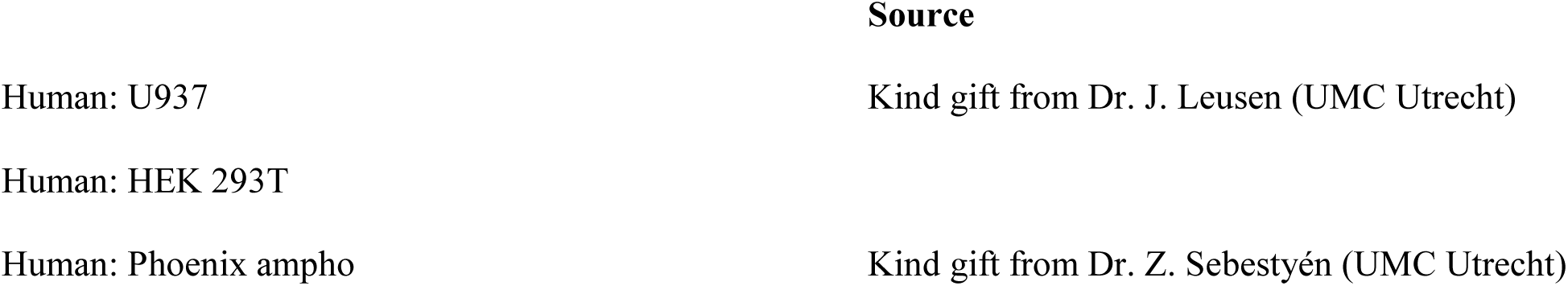

### Other

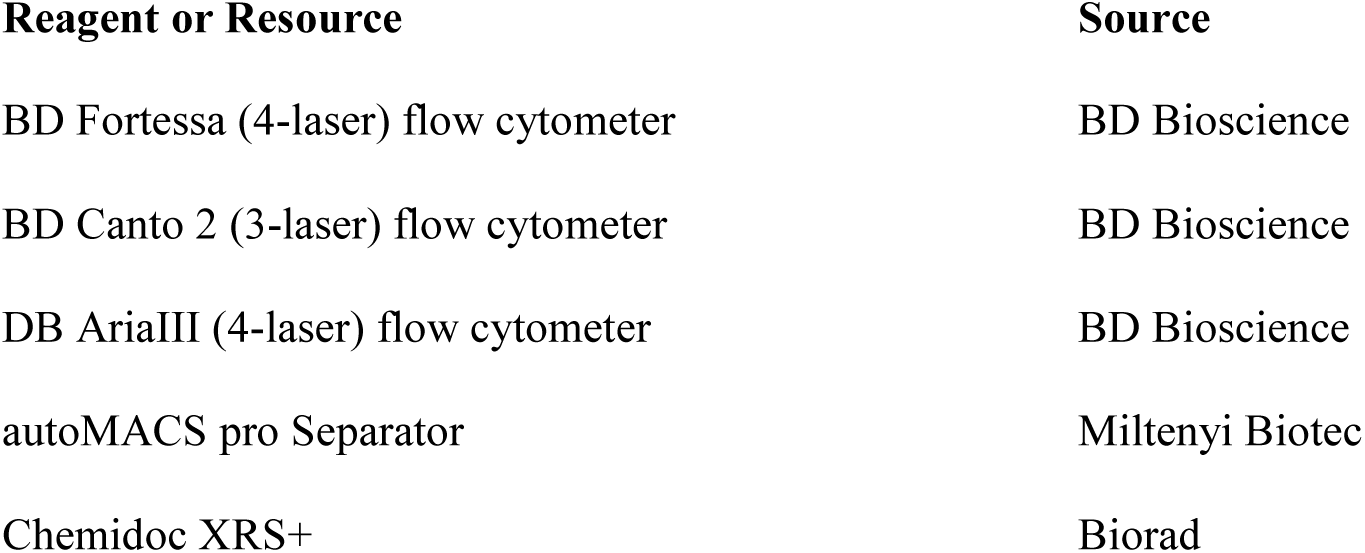

### Contact for Reagent and Resource Sharing

Further information and requests for resources and reagents should be directed to, and will be fulfilled by, Linde Meyaard. All data is available in the manuscript or the supplementary materials. Some materials used in this manuscript are subject to Material Transfer Agreement (MTA), or can only be distributed under MTA.

### Cells and cell culture

#### Human samples

SLE patients and age- and sex-matched healthy controls (HCs) were recruited from our outpatient clinic or in-house healthy donor service. None of the patients had evidence of an ongoing infection. The University Medical Center Utrecht medical ethical committee approved this study and all study participants signed informed consent forms. Please see Supplemental Table 1 for more detailed information on the human subjects in this study. PBMC from HC or SLE patients were isolated by ficoll gradient. Monocytes were isolated from PBMC with human CD14 MicroBeads (Miltenyi Biotec) MACS isolation using manual separation with LD columns (Miltenyi Biotec), or automated separation with an autoMACS pro Separator (Miltenyi Biotec). HC that were tested in functional experiments without direct comparison to SLE patients were all women in the age range of 18 to 62 years.

For moDC differentiation, monocytes were isolated from Buffy coats by ficoll gradient followed by percoll gradient and additional adhesion isolation. We are not informed about the sex and age of these donors. Monocytes were differentiated with GM-CSF (800U/ml) and IL-4 (500U/ml) in enriched RPMI (Gibco, 10% bovine serum, P/S, 2mM Glutamine) in a 5% CO_2_ humidified 37°C incubator as previously described (Geijtenbeek et al., 2003), and used at day 6 or 7 after the start of differentiation.

All cells were maintained at 37°C with 5% CO2 in a humidified cell culture incubator.

### Cell lines

U937 (human, male origin, not authenticated) cells were maintained in RPMI (Gibco) with 10% bovine serum and 1% P/S, 5% CO_2_ humidified 37°C incubator. Before experiments, U937 cells were differentiated with 30ng/ml Phorbol 12-myristate 13-acetate (PMA; Sigma, P8139-1MG) in 5% FCS RPMI for 24 hours, followed by a 24-hour rest period RPMI with 5% FCS. Unless otherwise indicated, U937 experiments were performed in 5% FCS containing medium.

CD200R-GFP was generated by adding GFP to the c-terminal domain of CD200R, preceded by a short flexible glycine-serine (GSGSGS) spacer. CD200RΔ-GFP was truncated directly after the predicted transmembrane domain. To obtain CD200R-GFP and CD200RΔ-GFP, U937 cells were transduced using standard retroviral transduction protocols and we used fluorescence-activated cell sorting to obtain equal expression levels.

U937 expressing full length CD200R-GFP were transduced with an in-house generated lentiviral construct containing both the gRNA and Cas9 (van de Weijer et al.). After transduction, cells were selected with puromycin and kept as poly-clonal lines. Successful deletion of the target gene was confirmed by intra-cellular antibody-staining followed by flow cytometry.

To overexpress RasGAP_1-455_ CD200R-GFP expressing cells were transduced with RasGAP-HA AA1-455.lti. After transduction, successful transduction was validated by intra-cellular flow cytometry.

### Experimental details

#### Stimulation assays: Human primary cells

For CD200R-ligation experiments, 10μg/ml culture grade anti-CD200R or isotype control; or 3μg/ml CD200his and CD200Rhis (both Sino Biologicals) were coated onto MaxiSORB (NUNC) plates at 4°C overnight in PBS.

Where indicated, PBMC or moDC were pre-incubated for 16 hours with 1000U/ml IFNα 2a (Cell Sciences) or enriched RPMI (RPMI, 10% bovine serum, P/S, 2mM Glutamine) in a 5% CO_2_ humidified 37°C incubator and washed with medium before proceeding with the experiment. Before seeding cells, the plates were washed three times with 100μl PBS. 100μl cell-suspension of moDC were seeded at 0.5E6 cells/ml and PBMC were seeded at 1E6/ml in enriched RPMI in a 96-well plate. After 2 hours of pre-incubation on the coated wells, we stimulated cells with 1 μg/ml of R848 (Invivogen). mRNA was isolated 4 hours after stimulation with R848, cell free supernatant was harvested 6 hours after stimulation with R848, and for flow cytometric analysis of phosphorylated proteins cells were fixed 30 minutes after R848 stimulation (see “PhosFlow Cytometry” for details).

#### Stimulation assays: U937-CD200R cells

PMA differentiated U937 cells were seeded in round bottom plates and incubated for 5 minutes at room temperature with anti-CD200R (in-house, OX108), anti-SIRL-1 (in-house, 1A5) as isotype control, CD200his or CD200Rhis (both Sino Biologicals) at 3μg/ml. For phosphorylation assays, cells were incubated with antibodies or recombinant proteins for 60 minutes in a 5% CO_2_ humidified 37°C incubator before fixation (see “PhosFlow” for details). For IL-8 secretion, cells were stimulated with 200ng/ml LPS (Salmonella Typhosa, Sigma) and cultured for 24 hours in a 5% CO_2_ humidified 37°C incubator before harvesting cell-free supernatant for ELISA.

#### Viral plasmids

CD200R was fused to eGFP by two-step sewing PCR. First CD200R or CD200RΔ, and eGFP were amplified with the following primers and appropriate sized fragments were isolated from agarose gels with the PCR clean-up Gel extraction (Macherey-Nagel). PCR fragments were mixed and a consecutive PCR was done with the CD200R-F and GFP-R primers before digestion with PmlI and NotI and ligation into pMX. We used sanger sequencing to validate the sequence in the plasmid matched the expected sequence.

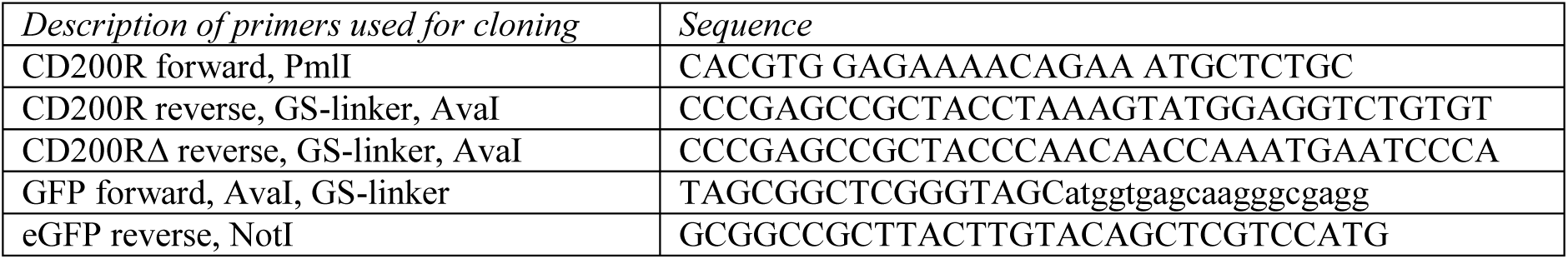

#### Viral transductions

HEK293T (human, female origin) were co-transfected with pMX-CD200R or CD200RΔ and pCL-Ampho to produce retroviral particles, RasGAP-lentiviral vectors together with pMDL, pVSV and pRSV. U937 were infected twice with cell-free virus supernatant in presence of 4ug/ml polybrene. After 5 days of culture, GFP^+^ U937 cells were sorted and selected for equal expression.

#### CRISPR/Cas9

The following sequences were cloned in RP-418 Cas9-puro:

gRNAs targeting DOK2:

#1: GCAGCAGCAGACGTTTGGAA

#2: GAGACGGGGCAGTGAAACA

gRNAs targeting RASA1:

#1: GATAGCAGAAGAACGCCTC

#2: GCAGAAGAACGCCTCAGGC

#### Western blot

For western blot analysis of RasGAP, monocytes or PBMC were lysed in Laemmli at a concentration of 1E7 cells/mL. Where indicated, PBMC or monocytes were pre-incubated for 16 hours with 1000U/ml IFNα 2a (Cell Sciences) before lysis. Lysates were homogenized by passing them three times though a 24G needle. Samples were loaded under denaturing conditions on pre-cast gels (Mini-PROTEAN TGX Gels, Any kD, Bio-Rad Laboratories), or home-made 10% acrylamide gels. Protein was transferred to PVDF membranes (Immobilon-P PVDF .45um, Merck Chemicals BV). Membranes were blocked in 5% fat-free milk (Elk, Campina, The Netherlands) in TBS 0.05% Tween-20 (TTBS), and incubated with antibodies in 1% Elk TTBS. The blots were imaged using SuperSignal West Femto (Thermo Fisher Scientific) on a Bio-rad Chemidoc MP.

#### mRNA isolation, cDNA synthesis and qPCR analysis

mRNA from the PBMC discovery cohort was isolated using an mRNA capture kit (Roche) and cDNA was synthesized with the Reverse transcriptase kit (Promega) according to manufacturer protocol. mRNA from all other PBMC cohorts was isolated using QIAGEN mRNA micro isolation kits, followed by iScript (Biorad) cDNA synthesis according to manufacturer protocol. qPCR was performed on a QuantStudio 12K Flex or a StepOnePlus Realtime PCR system (AB Instruments) with SYBR Select Master Mix (Life Technologies), or with TaqMan, where indicated, with 5ng cDNA input and 400nM of each primer using the fast protocol with melt curve as is suggested for SYBR Select Master Mix.

Relative expression = 2^(Ct(*B2M*)-Ct(target)).

#### Flow cytometry

Cells were washed in flow cytometry buffer (PBS, 1% BSA, 0.01% sodium azide, from here on FACS buffer), and incubated with antibodies for 30 minutes on ice, washed twice and analyzed on a BD Fortessa or BD Canto II using DIVA 8 software. For multi-color experiments, compensation settings were generated using the compensation tool in DIVA 8. For data analysis we used FlowJo V10 to gate for live cells, single cells and subsequent gating as indicated in supporting figures.

#### Intra cellular flow cytometry for phosphorylated proteins (PhosFlow)

For analyses of phosphorylated proteins with flow cytometry, cells were fixed after stimulations with a final concentration of 3% PFA in medium for 10 minutes at room temperature. Attached cells were detached by scraping with a plunger. Cells were harvested, transferred into a 96-well V-bottom plate and washed, and subsequently permeabilized with 100% ice old methanol for at least 5 minutes and a maximum of 2 weeks at −20°C.

Before continuing with the staining protocol, the PFA/Methanol fixed cells were washed twice. Staining was performed in flow cytometry buffer and non-specific binding was prevented by incubating the cells with 5% normal goat or mouse serum for 45 minutes at room temperature. The primary antibodies were incubated shaking for 1 hour at room temperature or 16 hours at 4°C. Cells were washed three times in 150μl FACS buffer and incubated with a conjugated secondary antibody for 1 hour at room temperature while shaking. After washing three times, fluorescence was assessed on a BD Fortessa or BD Canto II using a high throughput sampler attached to the flow cytometers.

‘live’ single cells were gated and median fluorescent intensity was used for quantification. For U937-CD200R and –CD200RΔ cells, cells were gated for similar expression of CD200R-GFP before assessing phosphorylation staining.

#### ELISA

ELISA kits used were all from Life Technologies: IL-8 (88-8086-88); IL-10 (88-7106-88); TNF (88-7346-88); IL-1β (88-7261-88) and IFNγ (88-7316-88). Cell-free supernatant was harvested and stored at −20°C until assayed by ELISA according to the manufacturer’s protocol. MAXIsorb plates (NUNC) were coated with 50μL capture antibody diluted in coating buffer for 16h at 4C. The plates were washed once 200 μL with PBC 0.05% Tween-20, and blocked with 100μL blocking buffer provided with the kit. The standard was diluted as the manufacturer recommended, and both the standard curve samples and assay samples were incubated for 2h at room temperature or 16h at 4°C while shaking. Plates were washed 3 times with 200μL PBS 0.05% Tween-20, and incubated 1 hour at room temperature with 50μL detection antibody in blocking buffer, followed by 3 times 200μL washes and incubation with strep-HRP in blocking buffer. Before development with TMB substrate, the plates were washed 5 times with 200μL PBS 0.05% Tween-20. Optical densities (OD) were measured at 450nM and 560nM as control. With PRISM 7.1 we extrapolated from a 4-parameter dose response curve based on the concentrations of the standard curve and their OD-values.

#### Calculations

We calculated the percentage inhibition as follows: (1-(target^anti-CD200R^/target^isotype^))*100. and categorized the effect of CD200R as follows: *neutral*: less than 15% inhibition or potentiation; *inhibition*: ≥ 15% inhibition; *potentiation*: ≥15% potentiation. We tested the effect of other thresholding ranging from 5% to 50% inhibition and potentiation, which did not affect the overall conclusion (not shown).The ratio to depict CD200R function was calculated as: *target*^anti-CD200R^/*target*^*i*sotype^ expression so that a value below 1 represents inhibition, and a value above 1 potentiation.

The inhibitory capacity of CD200R was calculated as follows: (% inhibition in the condition of interest)/ (% inhibition in CD200R-GFP U937) x 100. A value of 100 indicates that the treatment did not affect CD200R function and a value of 0 indicates complete loss of function.

#### Statistics

In all figure legends we have indicated the statistical test used to determine significance, and the n of experiments, or the data of individual donors is depicted. Statistical analysis was performed in Prism 7.1 (GraphPad software) or in R 5.3.1. All tests in this study are two-sided. All boxplots are plotted as median with 25 and 75 percentiles with handles range from minimum to maximum values. Bars are mean±SEM.

Significance is depicted as: ns: p<.1; *: p<.05; **: p<0.01; ***: p<0.001; ****: p<0.0001

