## Supplementary material for "Inhibitory CD200-receptor signaling is rewired by type I interferon": Fig. S

Vlist et al. Supplemental Figure 1

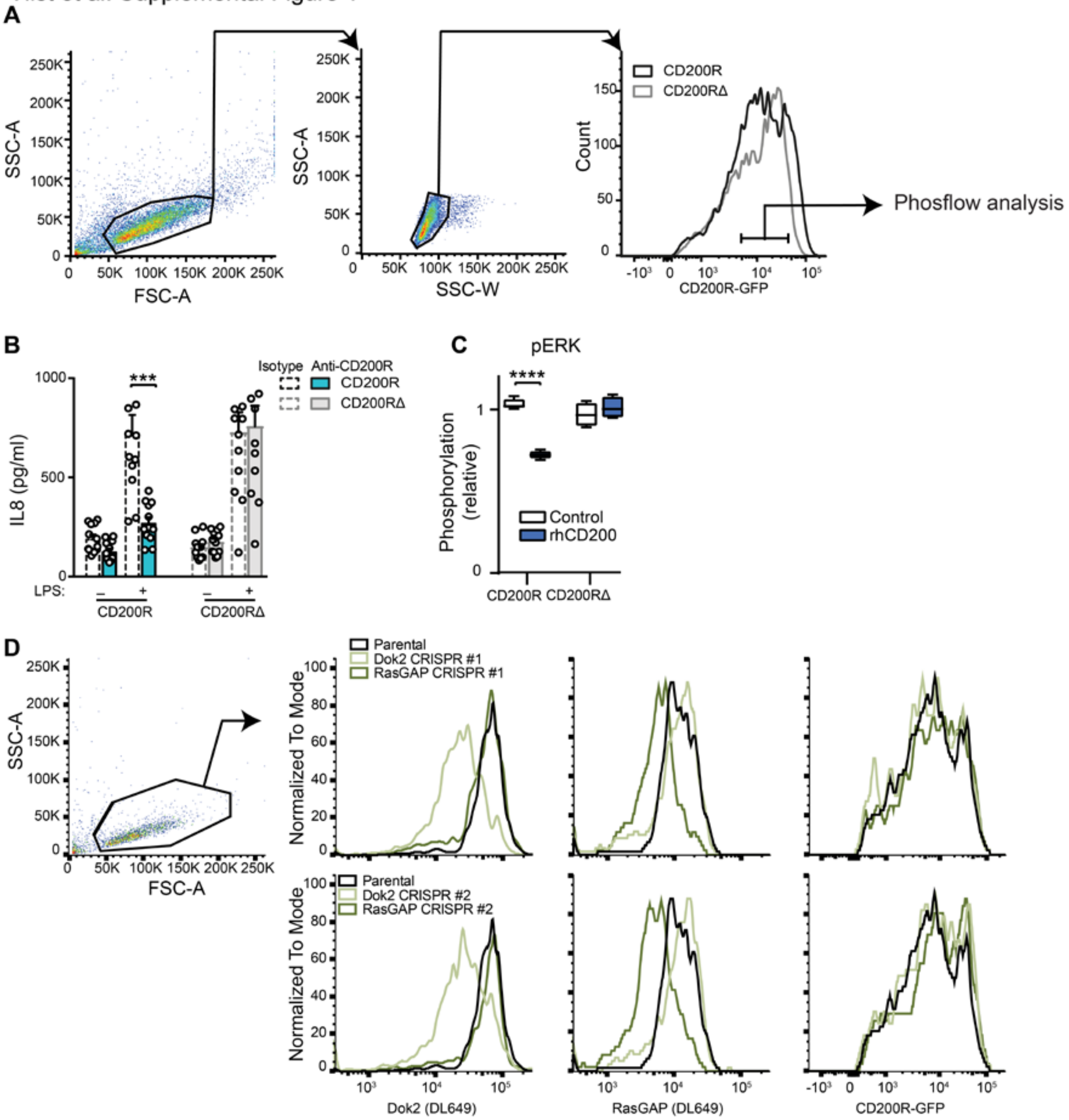

Supplemental Figure 1 – expression of CD200R adaptor molecules and function of CD200R.

(**A**) Gating strategy and expression of CD200R-GFP and CD200RΔ-GFP on U937 cells.

(**B**) IL-8 secretion by CD200R-U937 (blue) or CD200RΔ-U937 (gray) cells stimulated with LPS in the presence of agonistic anti-CD200R (solid line) or isotype control (dashed line). Data are shown as mean ± SEM. Significance was determined by a paired t test, n=12 from 6 experiments.

(**C**) Relative p-Erk in CD200R-U937 or CD200RΔ-U937 after rhCD200R (control, open) or rhCD200 (blue) stimulation. N=4 from two experiments. Significance was determined by a Welch's t test. Box represents median with 25 and 75 percentiles and bars represent the minimum to maximum values.

(**D**) Intracellular flow cytometric analysis of Dok2, RasGAP and CD200R-GFP expression in two polyclonal CD200R-U937 lines in which DOK2 was targeted, and two polyclonal lines in which RASA1 was targeted by CRISPR/Cas9.

Vlist et al. Supplemental Figure 2

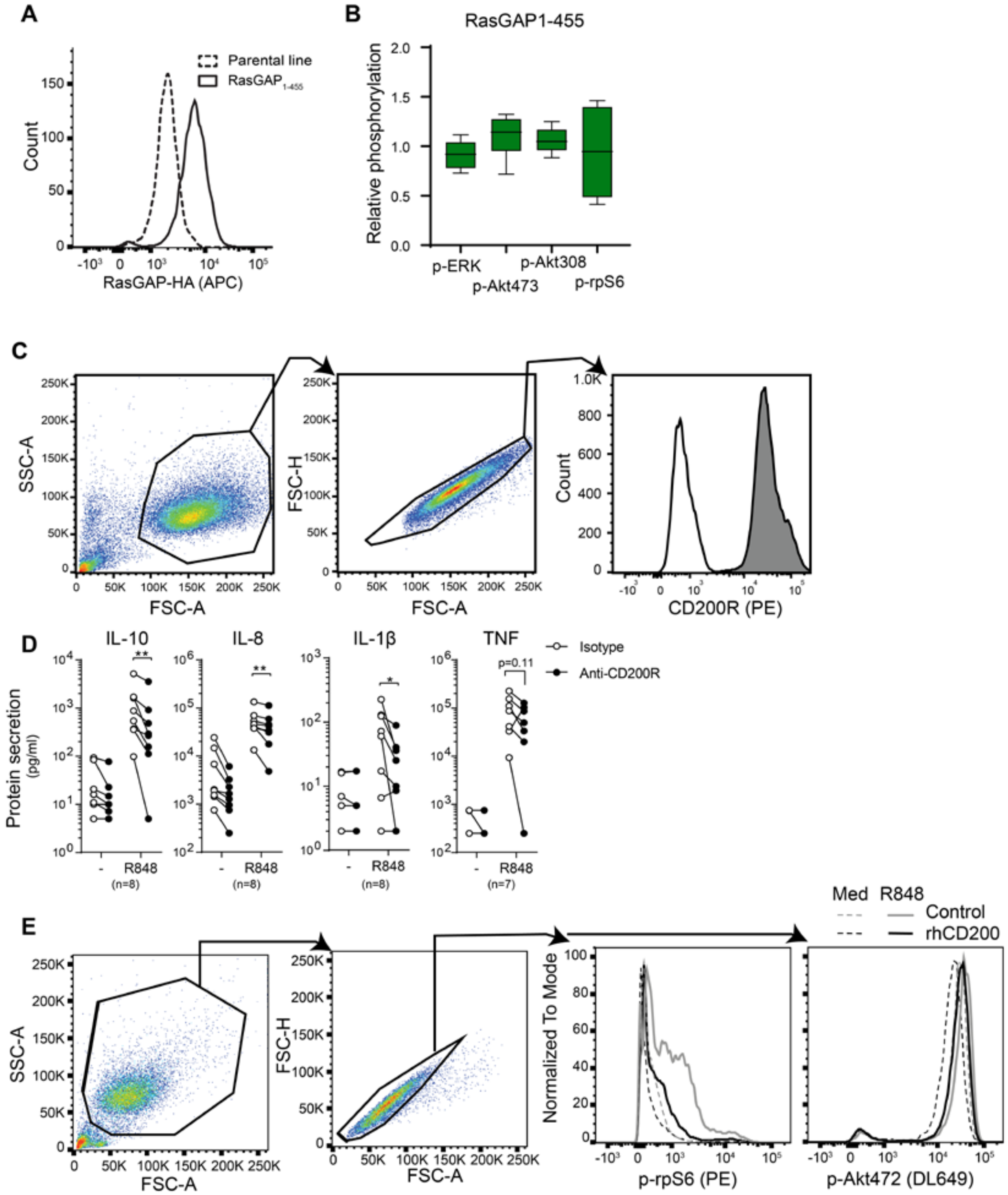

Supplemental figure 2 – CD200R signaling in primary cells.

(**A**) Expression of $RasGAP_{1\text{-}455}$-HA in U937 cells.

(**B**) The effect of overexpression of $RasGAP_{1\text{-}455}$ on basal phosphorylation of p-Akt308, p-Akt473, p-Erk, and p-S6 as determined by flow cytometry. Data are normalized to the phosphorylation in parental cells used in the same experiments. n=6. Significance was determined by one-sample T test.

(**C**) Gating strategy and expression of CD200R on human moDC as determined by flow cytometry.

(**D**) Cytokine release by moDCs stimulated with R848 while seeded on isotype control or anti-CD200R coated wells. Significance was determined by Wilcoxon signed-rank test, n=8 for IL-10, IL-8 and IL-1β, and n=7 for TNF.

(**E**) Gating strategy and example histograms of p-rpS6 and p-Akt473 in moDCs stimulated with R848 while coated on control (rhCD200R) or rhCD200.

Vlist et al. Supplmental Figure 3

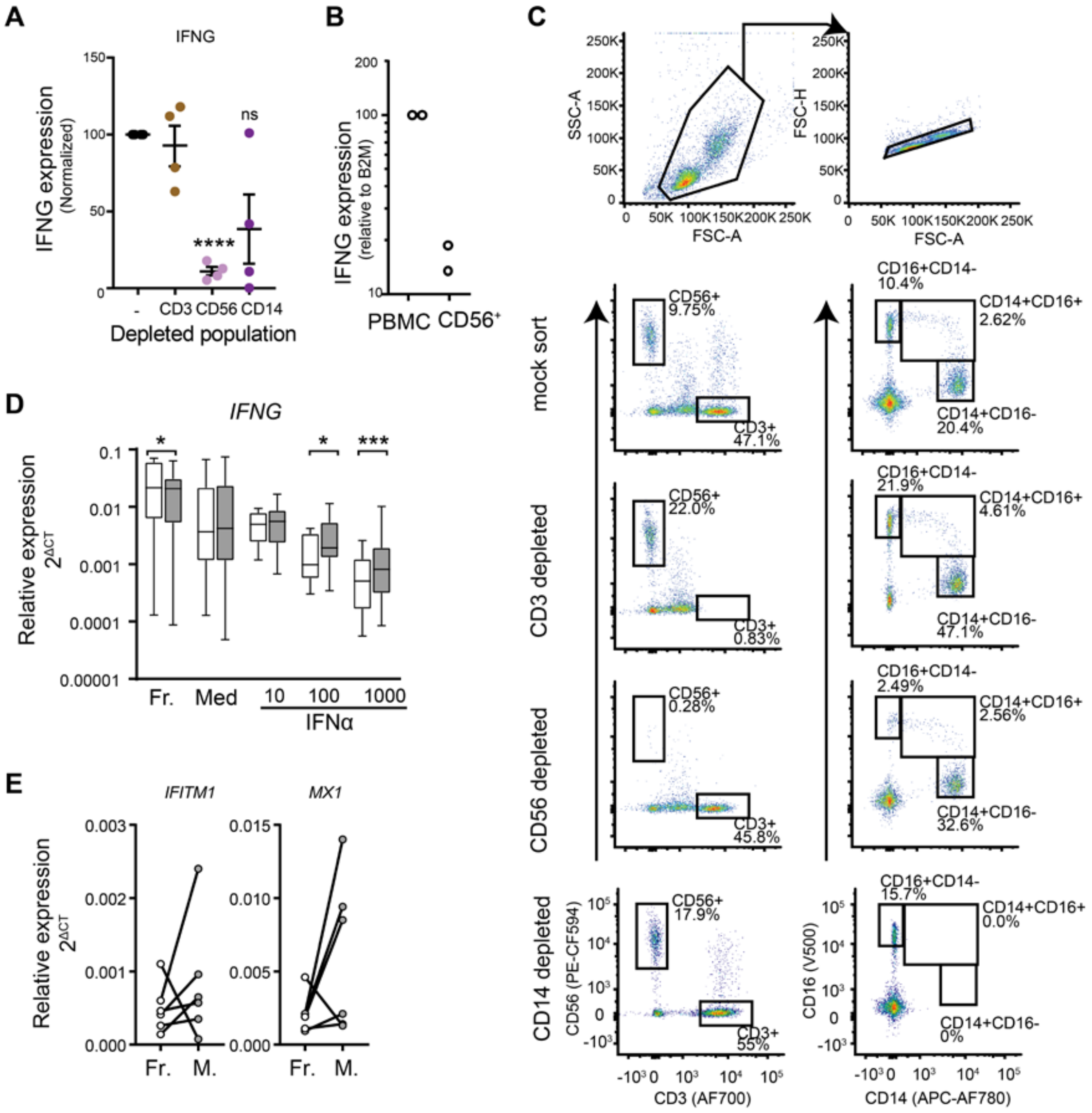


Supplemental figure 3.The effect of type I IFN on CD200R function.

(**A**) PBMC depleted for nothing (-), CD3 cells, CD56 cells or CD14 cells were stimulated with R848, and IFNG mRNA relative to B2M was normalized to the response in mock-sorted (-) PBMC. Significance was tested with a one-sample t test.

(**B**) IFNG mRNA production by NK cells 4 hours after stimulation with R848, normalized to mock-sorted PBMC.

(**C**) Gating strategy and scatter plots for expression of CD3, CD14 and CD56 of PBMC depleted for CD3, CD14 or CD56.

(**D**) Relative IFNG mRNA expression used to generate figure 3. *IFNG* mRNA production was measured in healthy control PBMC that were used fresh after isolation (Fr.), or have been cultured for 16h in medium (Med), 10, 100 or 1000U/ml of IFNα prior to ligation of CD200R with an agonistic antibody and stimulation of TLR7/8 with R848 for 4h. Significance was determined by a paired t test with Welch correction.

(**E**) Monocytes were CD14 MACS isolated and cultured for 16 hours. mRNA was isolated and expression of *IFITM1* and *MX1* was assessed relative to *B2M*. Fr.: Fresh. M.: Cultured for 16 hours in medium. Data represent 6 donors in three independent experiments.

Vlist et al. Supplemental Figure 4

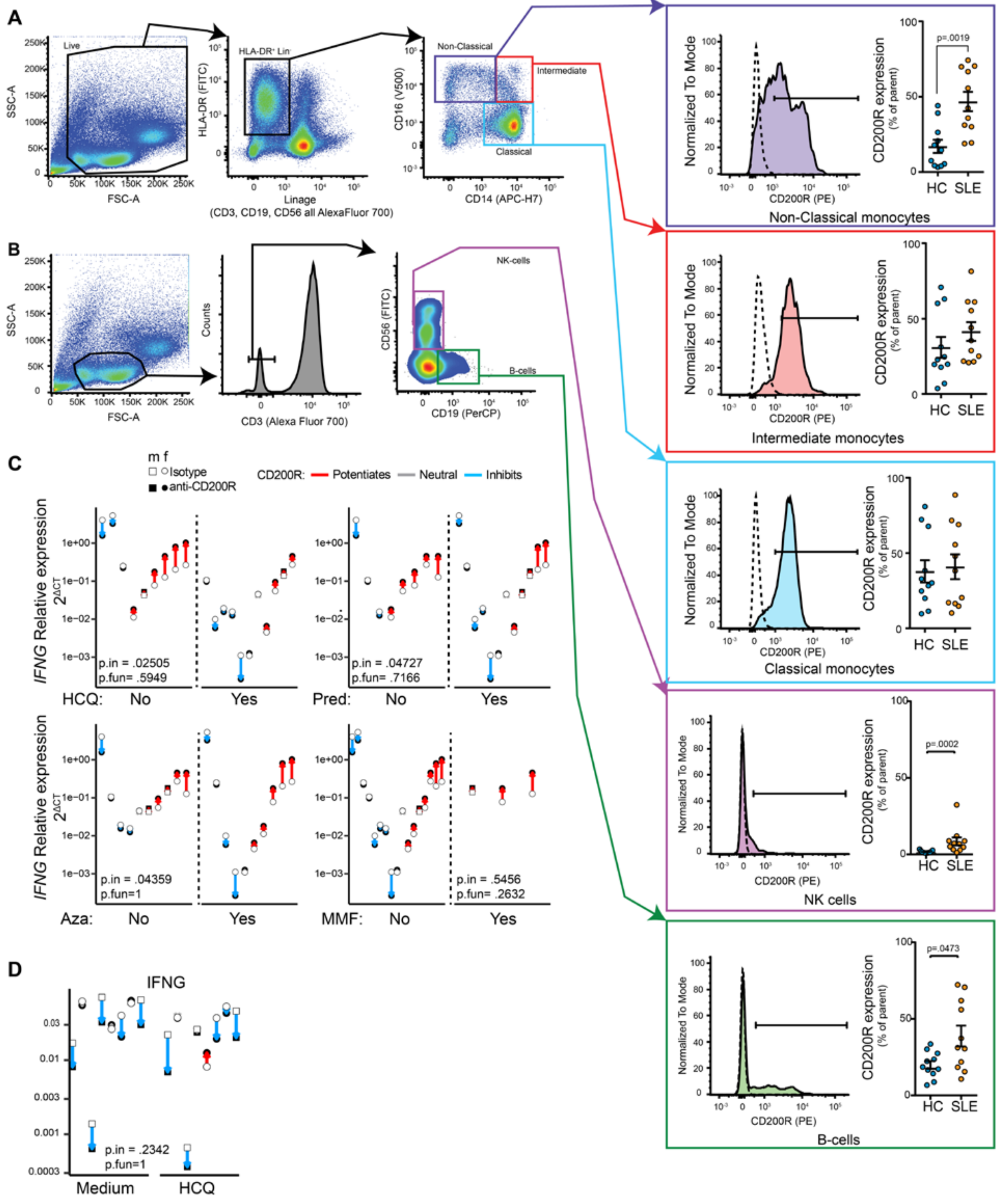

Supplemental figure 4 – CD200R function in systemic lupus erythematosus patients.

**(A, B)** Summarized expression data and gating strategy for monocytes, B cells and NK cells in frozen isolated PBMC from HC and SLE patients in our first cohort (Table S1). Graphs represent the percentage of CD200R of the parental gate. Filled histograms are anti-CD200R-PE stained samples and dashed lines are isotype control stained cells. See table S1. Significance was calculated with a Mann-Whitney test. Error bars represent mean±SEM.

(**C**) mRNA production of *IFNG* relative to *B2M* in PBMC stimulated with R848 in cohorts one and two (table S1) differentiated for use of hydroxychloroquine (HCQ), prednisone (Pred), azathioprine (Aza) and Mycophenolate mofetil (MFF). For effects of drugs on R848-induced *IFNG* mRNA production on PBMC seeded on isotype control (p.in) was tested by t test with welch correction. An effect of drugs on CD200R function (p.fun) was tested by Fishers Exact test, were we tested for the distribution of potentiating, neutral and inhibiting donors.

(**D**) The effect of HCQ on CD200R function and *IFNG* induction by R848 in healthy controls. Effects of HCQ on induction (p.in) was tested by t test with welch correction, and for an effect of HCQ on CD200R function (p.fun) was tested by Fishers Exact test.

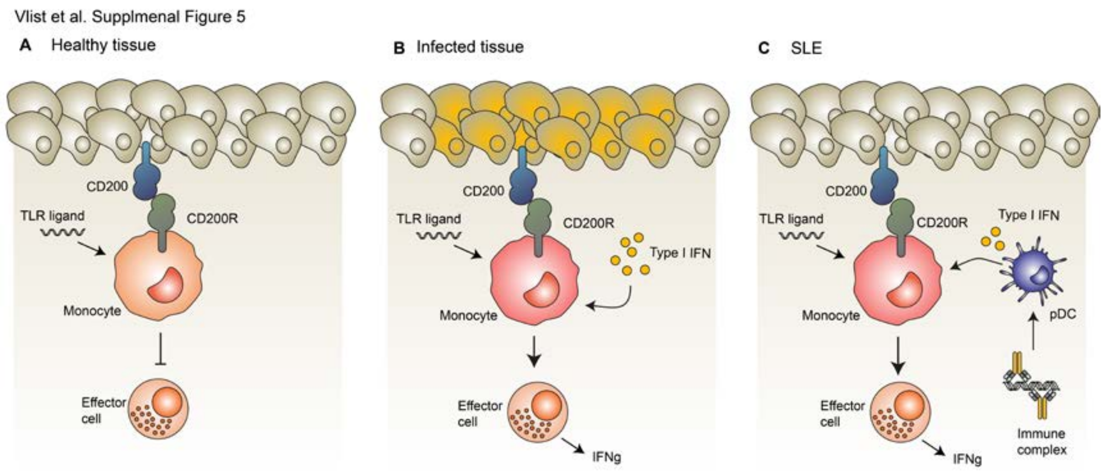


<u>Supplemental figure 5 – CD200R function health and disease.</u>

(**A**) In homeostasis, CD200-CD200R interaction suppresses immune responses because there is no need to respond vigorously to TLR ligand without context of inflammation.

(**B**) TLR ligands in presence of other inflammatory stimuli change CD200R into a potentiating receptor. We propose that this is beneficial in case of infections to feed-forward immune responses and help clear intracellular pathogens.

(**C**) Continuous exposure to auto-antibody-DNA/RNA complexes drives type I IFN production by pDCs. This augments immune activation in SLE patients and contributes to SLE pathogenesis.

| | Cohort 1 | | Cohort 2 | |
|---|---|---|---|---|
| | **SLE** | **Controls** | **SLE** | **Controls** |
| **N** | 11 | 11 | 11 | 14 |
| **Age (y, range)** | 39 (31-61) | 44 (31-67) | 47 (27–58) | 37 (26 – 58) |
| **Female (%)** | 100 | 100 | 82% | 79% |
| **SLEDAI (Mean±SD)** | 6 (4-6) | - | 4 (0- 4) | - |
| **C3 (g/L)** | 0.87 (0.72-1.01) | - | 0.98 (0.9-1.25) | - |
| **C4 (g/L)** | 0.14 (0.13-0.16) | - | 0.19 (0.13-0.25) | - |
| **Anti-dsDNA (IU/mL)** | 16 (5-128) | - | 2.5 (0.9 -17) | - |
| **Lupus Nephritis** | 64% | - | 55 % | - |
| **Hydroxychloroquine** | 18,2% | - | 82 % | - |
| **Prednisone** | 45.5% | - | 73 % | - |
| **MMF** | 18.2% | - | 9% | - |
| **Aza** | 45.5% | - | 45% | - |

Supplemental table 1. Cohort characteristics

Patient and healthy control description. Medians with interquartile range or percentage of total.
